# The role of dendritic brain-derived neurotrophic factor transcripts on altered inhibitory circuitry in depression

**DOI:** 10.1101/333294

**Authors:** Hyunjung Oh, Sean C. Piantadosi, Brad R. Rocco, David A. Lewis, Simon C. Watkins, Etienne Sibille

**Author notes:** Correspondence: E Sibille, CAMH, 250 College Street, Room 134, Toronto, ON M5T 1R8, Canada. Email address; xt36751.

## Abstract

**Background:** A parallel downregulation of brain-derived neurotrophic factor (*BDNF*) and somatostatin (*SST*), a marker of inhibitory γ-amino-butyric acid (GABA) interneurons which target pyramidal cell dendrites, has been reported in several brain areas of subjects with major depressive disorder (MDD), and rodent genetic studies suggests they are linked and both contribute to the illness. However, the mechanism by which they contribute to the pathophysiology of the illness has remained elusive.

**Methods:** With qPCR, we determined the expression level of *BDNF* transcript variants and synaptic markers in the prefrontal cortex (PFC) of MDD patients and matched controls (n=19/group) and of C57BL/6J mice exposed to chronic stress or control conditions (n=12/group). We next suppressed *BDNF* transcripts with long 3’ untranslated region (L-3’-UTR) using small hairpin RNA (shRNA) and investigated changes in cell morphology, gene expression and behavior.

**Results:** L-3’-UTR containing *BDNF* mRNAs, which migrate to distal dendrites of pyramidal neurons, are selectively reduced and highly correlated with *SST* expression in the PFC of MDD subjects. A similar downregulation occurs in mice submitted to chronic stress. We next show that *Bdnf* L-3’-UTR knockdown is sufficient to induce (i) dendritic shrinkage in cortical neurons, (ii) cell-specific MDD-like gene changes (including *Sst* downregulation), and (iii) depressive-/anxiety-like behaviors. The translational validity of the *Bdnf* L-3’-UTR shRNA-treated mice was confirmed by significant cross-species correlation of changes in MDD-associated gene expression.

**Conclusion:** These findings provide evidence for a novel MDD-related pathological mechanism linking local neurotrophic support, pyramidal cell structure, dendritic inhibition and mood regulation.

## INTRODUCTION

MDD is characterized by persistent low mood and/or anhedonia, and cognitive impairment. These symptoms are associated with dysfunction of brain networks involved in cognition, emotion and reward (1). The PFC plays a role in cognition and emotion regulation and is affected by stress (2). Consistent with functional impairment and atrophy (3-5), postmortem studies reported decreased neuronal body size, spine loss, and reduced cellular density in the PFC of MDD subjects (6,7).

Decreased GABAergic neurotransmission in MDD has been revealed by transcranial magnetic stimulation (8) and proton magnetic resonance spectroscopy (9,10). Decreased expression of GABA-related genes has been observed in corticolimbic areas of MDD patients (11-14). Downregulation of *SST*, a molecular marker of dendritic-targeting interneurons was consistently found, whereas parvalbumin (*PVALB*), the marker of perisomatic-targeting interneurons, remained relatively unaffected by disease, although see Tripp et al (14). Downregulation of *SST* is also observed in other neurological diseases (15), suggesting a vulnerability of GABA interneurons coexpressing this marker. The underlying mechanism is unknown, however, chronic stress and corticosterone exposure induced endoplasmic reticulum stress in these neurons (16). Another mechanism could be the loss of neurotrophic support. MDD and prolonged stress reduce *BDNF* expression (17,18), which might compromise structural integrity of various cells (19,20). GABAergic cells do not produce *BDNF* and rely on supply from other cell populations, mostly pyramidal neurons (21-23). In MDD, SST reduction is accompanied by *BDNF* and/or *NTRK2* (also known as *TRKB*) downregulation (11,14) and genetic studies in mice revealed that *SST* expression is highly dependent on *BDNF* (11,14,24), together suggesting a pathogenic mechanism linking *BDNF* and GABA.

Local action of *BDNF* is closely associated with neural plasticity. In addition to its autocrine action regulating dendritic and synaptic plasticity of pyramidal cells (25,26), *BDNF* can alter the presynaptic GABAergic system in a paracrine manner (23,27). In a previous transcriptome analysis of human PFC,

we found that the expression of *GABRA5* (gamma-aminobutyric acid A receptor, alpha 5), a subunit of GABA-A receptor enriched adjacent to dendritic GABAergic synapses and that mediates the function of *SST* neurons (28), showed the highest correlation with *BDNF* levels among all investigated genes (29). This suggests that local *BDNF* supply may play a role in maintaining function of *SST* neurons on pyramidal cell dendrites. Interestingly, *BDNF* transcripts are present in dendrites (30-32) and cellular localization of *BDNF* transcripts corresponds with phospho-*NTRK2* immunostaining (30), suggesting that *BDNF* may be locally translated and act on dendrites. Studies showed that dendritic *BDNF* is implicated in stress-related mood disorders: dendritic *BDNF* transcripts are decreased by stress (33-35), and increased by antidepressant treatment (36). Knockdown of dendritic *BDNF* impairs structural integrity of primary hippocampal neurons (30,37), replicating the effects of chronic stress.

Here, we investigated changes in dendritic *BDNF* transcripts in the dorsolateral PFC (dlPFC) of MDD subjects and medial PFC (mPFC) of mice exposed to chronic stress, and tested if such changes were linked to dendritic-targeting interneuron markers. Using shRNA, we then knocked-down dendritic *Bdnf* mRNA in primary neuronal culture and in mouse mPFC to assess whether reduced dendritic *Bdnf* expression was sufficient to induce MDD-and stress-related phenotypes. We predicted that decreased local neurotrophic support contributes to selective disturbance in dendritic structure and in dendritic-targeting GABAergic neurons, as a putative mechanism leading to mood dysregulation.

## METHODS AND MATERIALS

A complete description can be found in the Supplementary Information.

### Human postmortem subjects

Postmortem brains were collected during autopsies conducted at the Allegheny County Medical Examiner’s Office (Pittsburgh, PA) after consent from next of kin. For all cases, a committee of experienced clinicians makes consensus DSM-IV diagnoses using information obtained from clinical records, toxicology exam and standardized psychological autopsy (38). Individuals were screened for the absence of neurodegenerative disorders by neuropathological examination (39-41). All procedures were approved by the University of Pittsburgh’s Committee for the Oversight of Research and Clinical Trials Involving the Dead and Institutional Review Board for Biomedical Research. After careful examination of clinical and technical parameters, 19 pairs of MDD and unaffected control subjects were selected (Table S1).

### Animals and unpredictable chronic mild stress (UCMS)

9-10 week old C57BL/6J male mice were divided into two groups and submitted to control housing condition or UCMS consisting of a 7-week regimen of pseudo-random unpredictable mild stressors. The progression of the UCMS syndrome was monitored weekly by assessing coat state and weight changes for each mouse. All procedures were approved by the University of Pittsburgh Institutional Animal Care and Use Committee.

### Behavioral testing and emotionality z-score

Tests for depressive-and anxiety-like phenotypes included the Elevated Plus Maze (EPM), Open Field(OF), novelty suppressed feeding (NSF) and Cookie Tests (CT). To assess behavioral consistency across tests, we integrated emotionality-related measures across tests as described earlier (42).

### RNA extraction and real-time quantitative polymerase chain reaction (qPCR)

For human studies, gray matter of dlPFC were collected in Trizol and further purified with Qiagen RNeasy micro kit (QIAGEN, Valencia, CA, USA). For mice, mPFC punches were taken and processed with Qiagen RNeasy micro kit. To generate cDNA, total RNA was mixed with qScript cDNA supermix (Quanta BioSciences, Gaithersburg, MD, USA). PCR products were amplified in triplets on a Mastercycler real-time PCR machine (Eppendorf, Hamburg, Germany).

### shRNA, DNA constructs and Adeno-associated viral (AAV) vector

shRNA sequence targeting L-3’-UTR of mouse *Bdnf* was adapted from (37). Lentiviral vectors containing shRNA against *Bdnf* L-3’-UTR or scrambled shRNA, and eGFP reporter gene were purchased from GeneCopeia (Rockville, MD, USA). AAV9 vectors containing *Bdnf* L-3’-UTR shRNA-GFP or scrambled shRNA were commercially prepared (Virovek, Hayward, CA, USA).

### Primary culture, transfection and fluorescent-activated cell sorting (FACS) of mouse cortical neurons

Primary mouse cortical neurons (Thermo Fisher Scientific; Waltham, MA, USA; A15585) were cultured in Neurobasal media (Thermo Fisher Scientific; 21103) supplemented with B27 (Thermo Fisher Scientific; 17504), GlutaMAX™-I (Thermo Fisher Scientific; 35050). At 7 days in vitro (DIV7), transfection was performed using Lipofectamine 3000 (Thermo Fisher Scientific; L3000015). GFP-expressing cells were collected by FACS into Qiagen RNeasy lysis buffer. RNA extraction, cDNA synthesis and qPCR were performed as described above.

### Immunostaining and image analysis

After 5 days of transfection, cells were fixed with 4% paraformaldehyde (PFA) and stained with Alexa Fluor® 488 conjugated rabbit anti-GFP (Life Technologies, Carlsbad, CA, USA; A21311) and mouse anti-*Map2* (Sigma-aldrich, St. Louis, MO, USA; M9942) to visualize dendrites of transfected cells. The morphology of dendrites was automatically traced and analyzed by Imaris software (Bitplane AG, Concord, MA, USA).

### Animals and stereotaxic surgery

9-10 week old C57BL/6J male mice were bilaterally injected with AAV9-*Bdnf* L-3’-UTR shRNA-GFP or with AAV9-scrambled shRNA-GFP into the mPFC (A/P +2.0 mm, M/L ±0.4 mm, D/V −2.0 mm). Behavioral tests were performed before and after UCMS exposure. For gene expression analysis, mice received *Bdnf* L-3’-UTR shRNA in one hemisphere and scrambled shRNA in the other for intra-subject comparison.

### Fluorescent in situ hybridization (FISH) and laser capture microdissection (LCM)

AAV-infused mice were perfused and lightly fixed with 1 mL of 4% PFA. Mouse brains were sectioned and thaw-mounted onto a polyethylene naphthalate-membrane slides (Leica Microsystems, Concord, ON, Canada). *Sst*-and *Pvalb*-expressing neurons were visualized by FISH (RNAscope, Advanced Cell Diagnostics, Newark, CA, USA). Briefly, brain sections were permeabilized with protease and antisense probe targeting *Sst* and *Pvalb*, preamplifiers, amplifiers, and Atto 550 conjugated-probes were serially hybridized at 40°C. After dehydration, ∽200 *Sst*+ or *Pvalb*+ cells were collected using a LMD7 system (Leica Microsystems) and processed for RNA extraction.

### Statistical analysis

Gene expression differences between MDD and control subjects were determined by analysis of covariance (ANCOVA) using SPSS (SPSS, Inc., Chicago, IL, USA). To determine covariates to include in gene-specific models, nominal factors were tested as main factors by ANOVA, scale covariates were tested by Pearson correlation, and repeated measures were corrected by modified Holm-Bonferroni test. ANCOVA models including significant co-factors were then applied. For cell culture and animal studies, statistical significance between two groups was determined with Student’s t-test. Repeated measures ANOVA was performed to determine interaction between shRNA and stress, effect of stress or shRNA on coat state over time.

## RESULTS

### Parallel downregulation of dendritic *BDNF* transcripts and dendritic targeting interneuron markers in the PFC of human subjects with MDD

The *BDNF* gene is composed of at least nine exons and makes diverse transcript variants using a combination of alternative splicing and polyadenylation sites (Figure S1A). The spatial segregation of *BDNF* mRNA is encoded by different untranslated regions; transcript variants with exon 2, 6 and the L-3’-UTR can migrate to distal dendrites whereas mRNAs containing exon 1, 4 and the short 3’ UTR are restricted to the soma and proximal dendrites (30,43,44). We used qPCR to quantify *BDNF* expression, focusing on transcript variants with known cellular localization, its receptor *NTRK2*, and synaptic function-related genes whose expressions are closely linked to *BDNF* levels in human PFC (29). Primers targeting *BDNF* protein coding sequence (CDS) were included to measure pan-*BDNF* level. Several *BDNF* variants displayed nominal changes, but only L-3’-UTR was significantly reduced in MDD samples (F=6.98, p=0.012; Figure 1A). The dendritic localization of *BDNF* L-3’-UTR was confirmed in mouse primary cortical neurons and hippocampal tissue (Figure S1B-C). Expression changes were not observed for *NTRK2* isoforms and excitatory synaptic-related genes, however, inhibitory synaptic genes displayed disease-related downregulation (Figure 1B-E). Interestingly, genes for all three dendritic-targeting interneuron markers (*SST, NPY, CORT*) and for dendritically-localized *GABRA5* were decreased in MDD (*SST:* F=15.03, p=4.6E-04; *NPY:* F=10.56, p=0.003; *CORT:* F=19.76, p=8.5E-05; *GABRA5:* F=10.18, p=0.003), whereas expression of *PVALB*, the molecular marker of perisomatic-targeting interneurons, was not affected (F=1.94, p=0.173).

**Figure 1.**
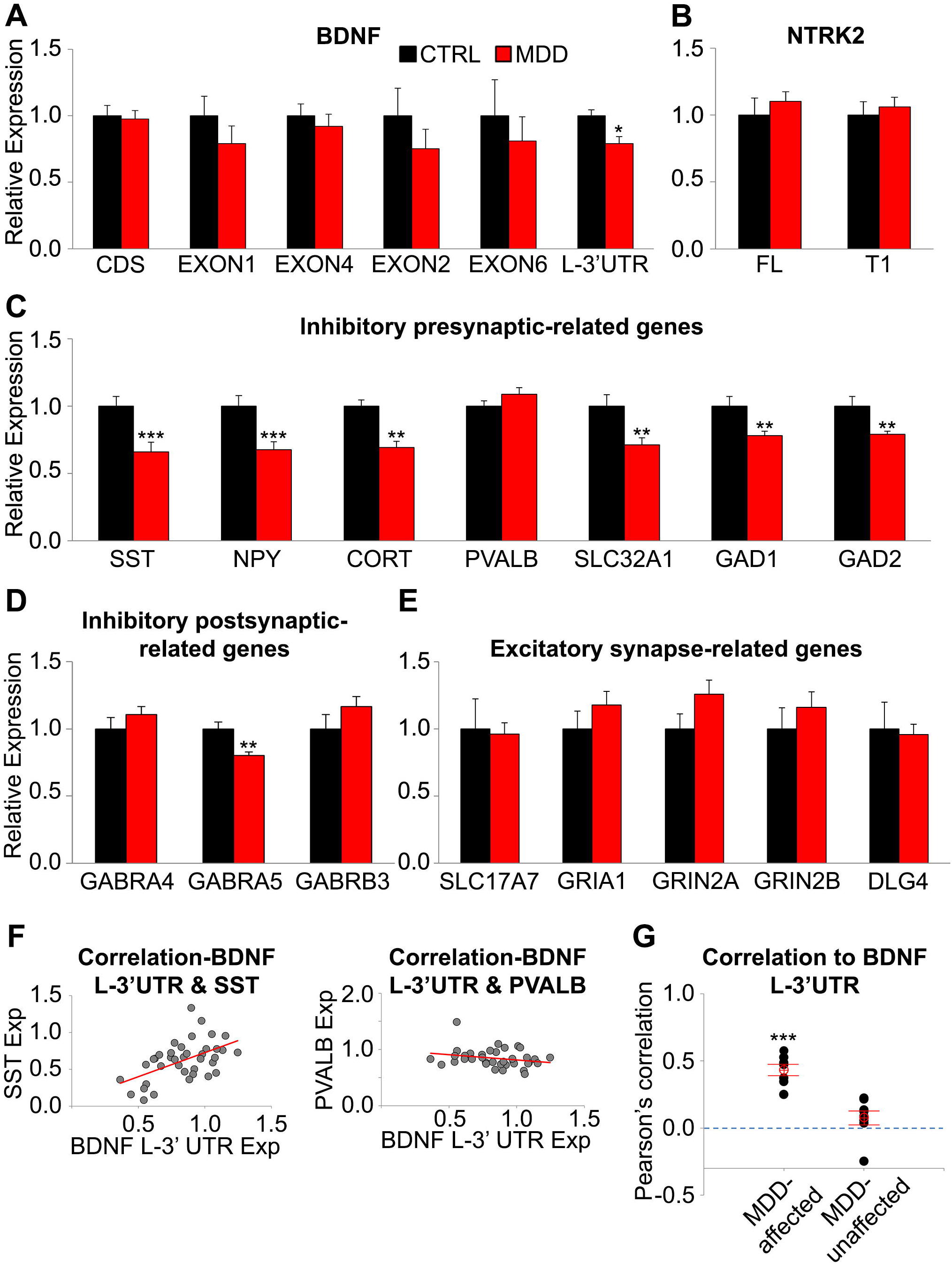
MDD-related changes in *BDNF* transcript variants and synaptic markers in human dlPFC. (A) Relative expression level (MDD/control) of *BDNF* transcripts, (B) *NTRK2* isoforms, and (C-E) selected synapse-related genes (n=19 subjects/group). BDNF EXON1 and 4 are associated with localization in soma and proximal dendrite. BDNF EXON2, 6 and L-3’-UTR are associated with localization in distal dendrites. (F) Pearson’s correlation between *BDNF* L-3’-UTR and *SST*, and *PVALB*. *BDNF* L-3’-UTR is positively correlated with *SST*, a dendritic targeting interneuron marker, but not with *PVALB*, a marker of perisomatic targeting interneurons. (G) Overall correlation between *BDNF* L-3’-UTR and MDD-affected or unaffected synaptic markers. Significant positive correlations between *BDNF* L-3’-UTR level and MDD-associated gene expression changes suggest a contribution of dendritic *BDNF* to transcriptome alterations. *, p<0.05; **, p<0.01; ***, p<0.001. Data are represented as mean ± SEM.

Despite the absence of *BDNF* messenger in interneurons (Figure S2), *BDNF* L-3’-UTR level showed positive correlations to MDD-affected genes such as *SST* (r=0.53, p=0.001), but not to genes unaffected by disease such as *PVALB* (r=-0.25, p=0.131; Figure 1F). The average correlation valuesbetween *BDNF* L-3’-UTR and MDD-affected or unaffected genes were 0.43±0.11 (p=5.7E-05) and 0.08±0.15 (p=0.178; Figure 1G), respectively. No other *BDNF* transcript variants including total *BDNF* showed significant correlationtoMDD-affectedgenes(CDS:r=-0.14±0.07, p=0.104;EXON1:r=-0.09±0.04, p=0.082; EXON4;r=-0.13±0.09, p=0.222; EXON2: r=0.11±0.10, p=0.310; EXON6: r=-0.09±0.09, p=0.320).

Consistent with previous finding (13), female MDD subjects showed greater *SST* reduction compared to male subjects (Table S2). However, correlation between BDNF-L-3’-UTR and SST was comparable in both sex (male: r=0.51, female: r=0.48). We observed age-related reduction of *BDNF* L-3’-UTR, *SST* and *CORT* transcripts in controls, however, such age-effects were less robust and not significant in MDD, consistent with prior findings in other cohorts (13,14). Other parameters including antidepressants or suicide were not consistently associated with the expression of all tested genes.

Together, selective downregulation in the expression of dendritic *BDNF* and dendritic-targeting interneuron markers was observed in the dlPFC of MDD subjects. The high correlation between expressions of *BDNF* L-3’-UTR and specific GABA-related genes suggests that reduced dendritic *BDNF* may be associated with functional alterations of dendritic-targeting GABAergic neurons.

### Chronic stress downregulates dendritic *Bdnf* transcript in the rodent mPFC

Next, we tested whether chronic stress, a major risk factor for MDD, was sufficient to induce MDD-like gene expression changes in mice. UCMS increased the latency to bite on day 2 of the cookie test (p=0.03; Figure 2A) and the food pellet in the NSF test (p=0.03; Figure 2B). The effect of stress was variable in other tests (Figure S3). To assess behavioral consistency across tests, we derived behavioral z-scores (16,42) and confirmed that UCMS-exposed mice exhibited higher depression-/anxiety-related behaviors, denoted as behavioral emotionality, compared to controls (p=0.003; Figure 2C).

**Figure 2.**
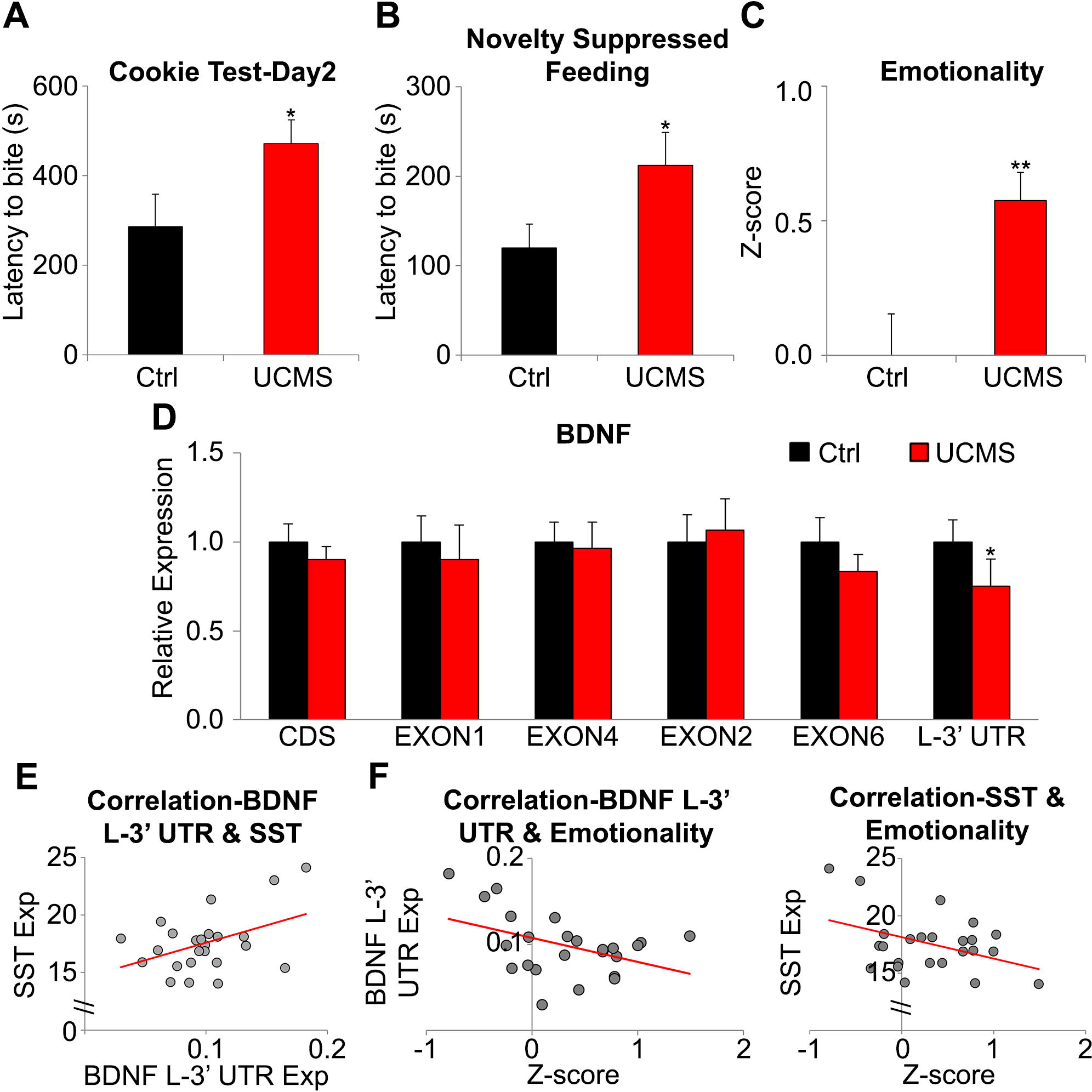
Chronic stress-related changes of anxiety-/depressive-like behaviors and expression of *Bdnf* transcript variants and *Sst* in mouse mPFC. (A) Stress-associated behavioral change in the cookie test (increased latency to bite cookie), (B) novelty suppressed feeding (increased latency to bite food pellet). (C) Increased depressive-/anxiety-like phenotype in stressed mice compared to controls. (D) Relative expression level (stress/control) of *Bdnf* transcript variants. (E) Positive correlation between *Bdnf* L-3’-UTR and *Sst* suggests a functional link between the two genes in mouse brain. (F) Behavioral emotionality was negatively correlated to *Bdnf* L-3’-UTR (left) and *Sst* (right). n=12/group;*p<0.05, **p<0.01. Data are represented as mean ± SEM.

We found a significant reduction of *Bdnf* L-3’-UTR in the mPFC of UCMS-exposed mice (p=0.029; Figure 2D). This expression change was not observed for other genes (Figure S4). Despite an absence of difference at the group level, there was a positive correlation between *Bdnf* L-3’-UTR and *Sst* (r=0.43, p=0.036; Figure 2E) and negative correlations between behavioral emotionality and both *Bdnf* L-3’-UTR (r=-0.43, p=0.036) and *Sst* (r=-0.41, p=0.047; Figure 2F). In comparison, *Pvalb* expression was not correlated with either *Bdnf* L-3’-UTR level (r=-0.23, p=0.281) or emotionality (r=0.17, p=0.420).

In summary, UCMS induced a selective downregulation in L-3’-UTR-containing *Bdnf* mRNAs in the mouse mPFC which was significantly correlated with behavioral emotionality and *Sst* levels. These results suggest a role for dendritic *Bdnf* in anxiety-/depressive-like measures via regulation of dendritic-targeting interneurons.

### Knockdown of dendritic *Bdnf* transcript is sufficient to induce MDD-and stress-associated phenotypes

We next tested whether altered dendritic *Bdnf* expression was causal to MDD/stress-related dendritic shrinkage of cortical neurons. Using GFP as a marker (Figure 3A), we collected shRNA-expressing primary neurons with FACS and confirmed that shRNA treatment knocked down *Bdnf* L-3’-UTR expression (Figure 3B), without affecting total *Bdnf* levels. shRNA expression reduced the number of intersections in distal dendrites without changes in proximal dendrites (0 to ∽30µm from soma) (Figure 3C, Table S3). The total length of all dendrites of shRNA-treated neurons was shorter than controls (p=0.009, Figure 3D) while the number of dendritic segments remained unchanged (p=0.91; Figure 3E).

**Figure 3.**
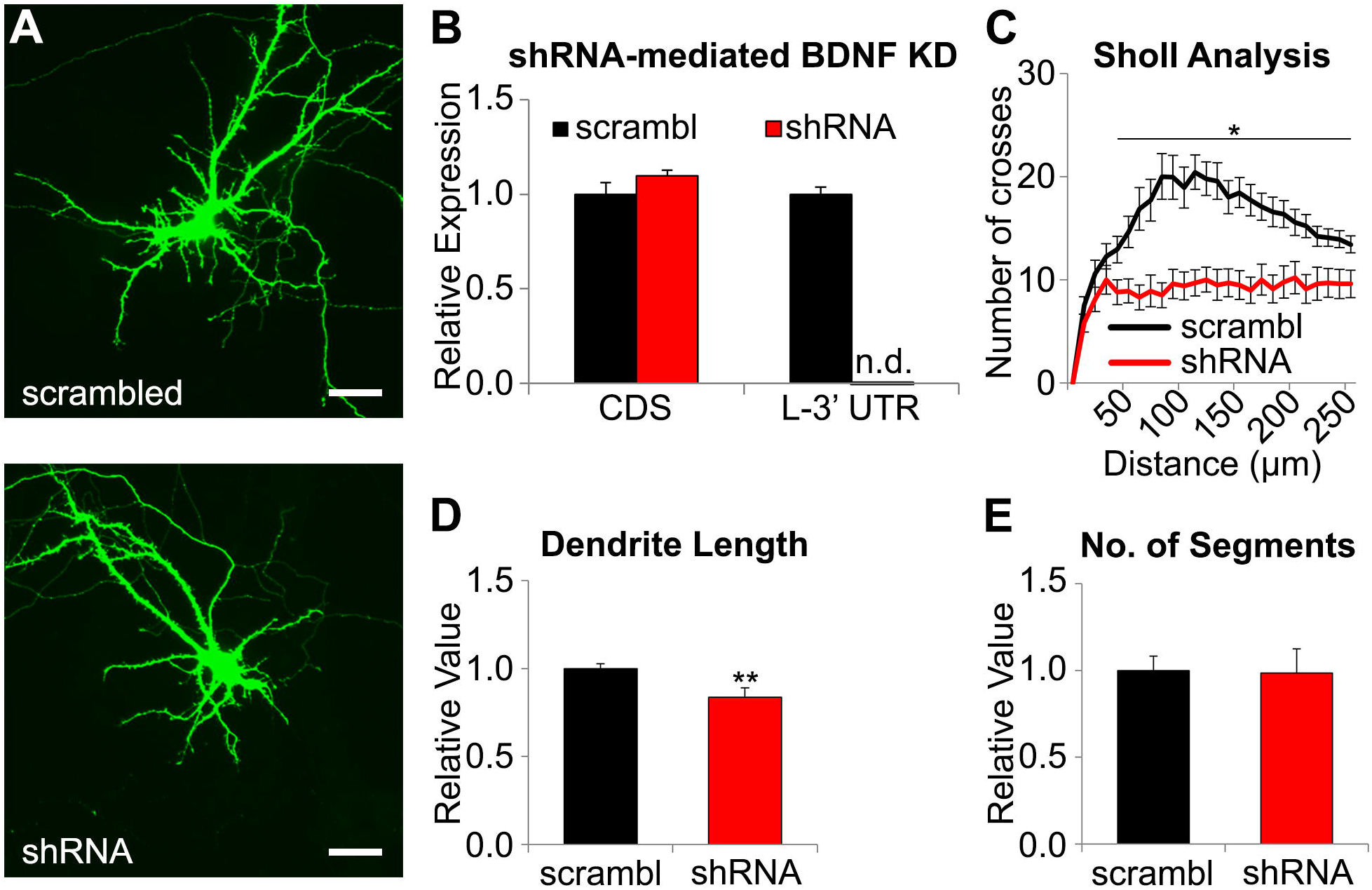
*Bdnf* L-3’-UTR knockdown-induced structural changes of primary mouse cortical neurons. (A) Representative image of primary mouse cortical neuron expressing scrambled shRNAand *Bdnf* L-3’-UTR-targeting shRNA. Scale bar = 50µm (B) Relative expression (shRNA/scrambled) of *Bdnf*-CDS and *Bdnf* L-3’-UTR in shRNA expressing cells compared to controls. Expression of L-3’-UTR was not detectable (n.d.) after shRNA treatment. (C) Sholl analysis for shRNA-treated and control neurons showed that shRNA treatment disrupted integrity of distal dendrite. (D) Relative value of total length of dendrites of shRNA-treated and control neurons. (E) Relative value of number of dendritic segments of shRNA-expressing neurons compared to control. Total dendritic length was significantly decreased without changes in branch number (control: n=14, shRNA: n=10; *p<0.05, **p<0.01). Data are represented as mean ± SEM.

Next, we injected shRNA-expressing AAV in mouse mPFC (Figure 4A) to investigate whether low dendritic *Bdnf* expression is associated with behavioral changes. Behavioral tests were performed after recovery, then after UCMS (Figure 4B). Stress exposure did not affect body weight (Figure 4C), but induced a progressive worsening of the coat state (Figure 4D), indicating that both groups responded to UCMS (p=3.6E-28). Notably, shRNA-treated mice displayed worse coat state than the control group over time (p=0.002). shRNA-treated mice showed trend-level differences in the OF (p=0.052; Figure 4E) and in emotionality z-scores before UCMS (p=0.093; Figure 4F).

**Figure 4.**
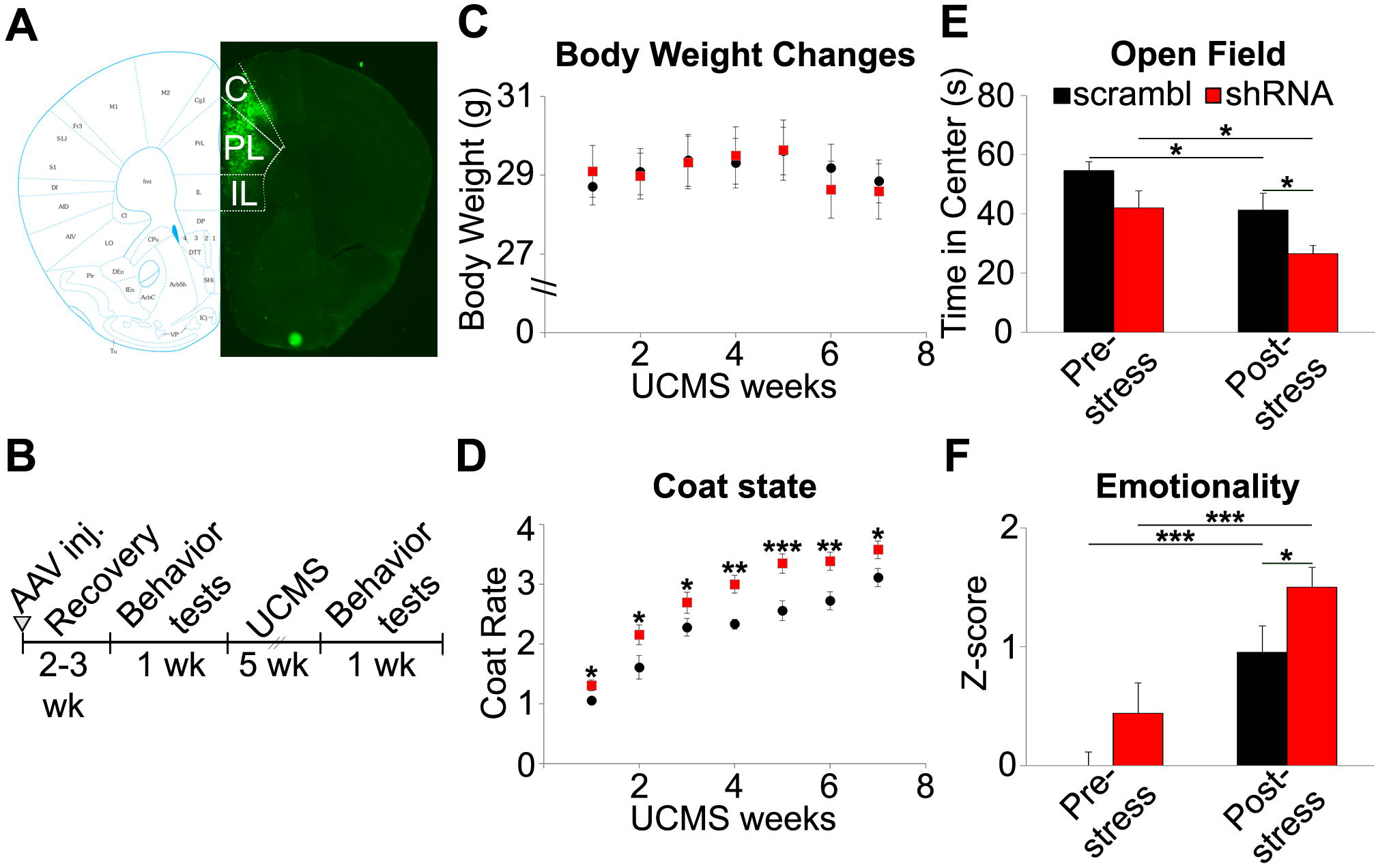
Effect of *Bdnf* L-3’-UTR knockdown and chronic stress on behavior. (A) Representative image of GFP expression following AAV injection in mPFC. The mouse mPFC map is from Paxinos and Franklin’s the mouse brain in stereotaxic coordinates (66). C: Cingulate cortex, PL: Prelimbic cortex, IL: Infralimbic cortex). (B) Schematic of the experimental design. (C) No body weight changes by chronic stress. (D) Progressive coat state changes with stress exposure. (E) shRNA-treated mice spent less time in the center of the open field apparatus after UCMS. (F) shRNA-treated mice group showed significantly higher emotionality z-scores than control after stress exposure (scrambled: n=9, shRNA: n=13; *p<0.05, **p<0.01, ***p<0.001). Data are represented as mean ± SEM.

After UCMS, shRNA-treated mice spent less time in the OF center (p=0.009; Figure 4E). Although other behavioral measures did not show significant group differences (Figure S5), shRNA-treated mice displayed higher emotionality compared to control vector-treated mice after UCMS (p=0.029; Figure 4F). There was no interaction between shRNA treatment and chronic stress on emotionality (p=0.763) or coat state (p=0.270).

To examine whether knockdown of dendritic *Bdnf* was sufficient to induce MDD-associated molecular changes, we infused AAV expressing *Bdnf* L-3’-UTR shRNA into one hemisphere and scrambled shRNA into the other (Figure 5A) and sacrificed mice at 2, 4, and 6 weeks after surgery. Gene expression changes were not found at post-surgery week 2. *Bdnf* L-3’-UTR was significantly downregulated at week 4 (p=0.013) and week 6 (p=0.030), whereas *Sst* level was reduced at week 6 only (p=0.002; Figure 5B). *Sst* and *Bdnf* L-3’-UTR expression showed positive correlation at week 6 (r=0.67, p=0.016), whereas *Pvalb* expression did not show such relationship (r=-0.05, p=0.866; Figure 5C), confirming the selective effect of *Bdnf* L-3’-UTR on *Sst*. In contrast to the knockdown yields observed in primary culture, total and EXON4+ *Bdnf* levels were reduced by shRNA (p=0.009 and 0.035, respectively; Figure S6A), suggesting differences between *in vitro* and *in vivo Bdnf* transcripts regulation, and/or *in vivo* functional adaptations. Notably, total and EXON4+ *Bdnf* levels were not correlated with *Sst* or *Pvalb* levels (total to *Sst:* r=0.18, p=0.566, to *Pvalb:* r=-0.15, p=0.646; EXON4 to *Sst:* r=0.39, p=0.205, to *Pvalb:* r=0.12, p=0.716). The expression of other genes was not changed by shRNA treatment (Figure S6B-E).

**Figure 5.**
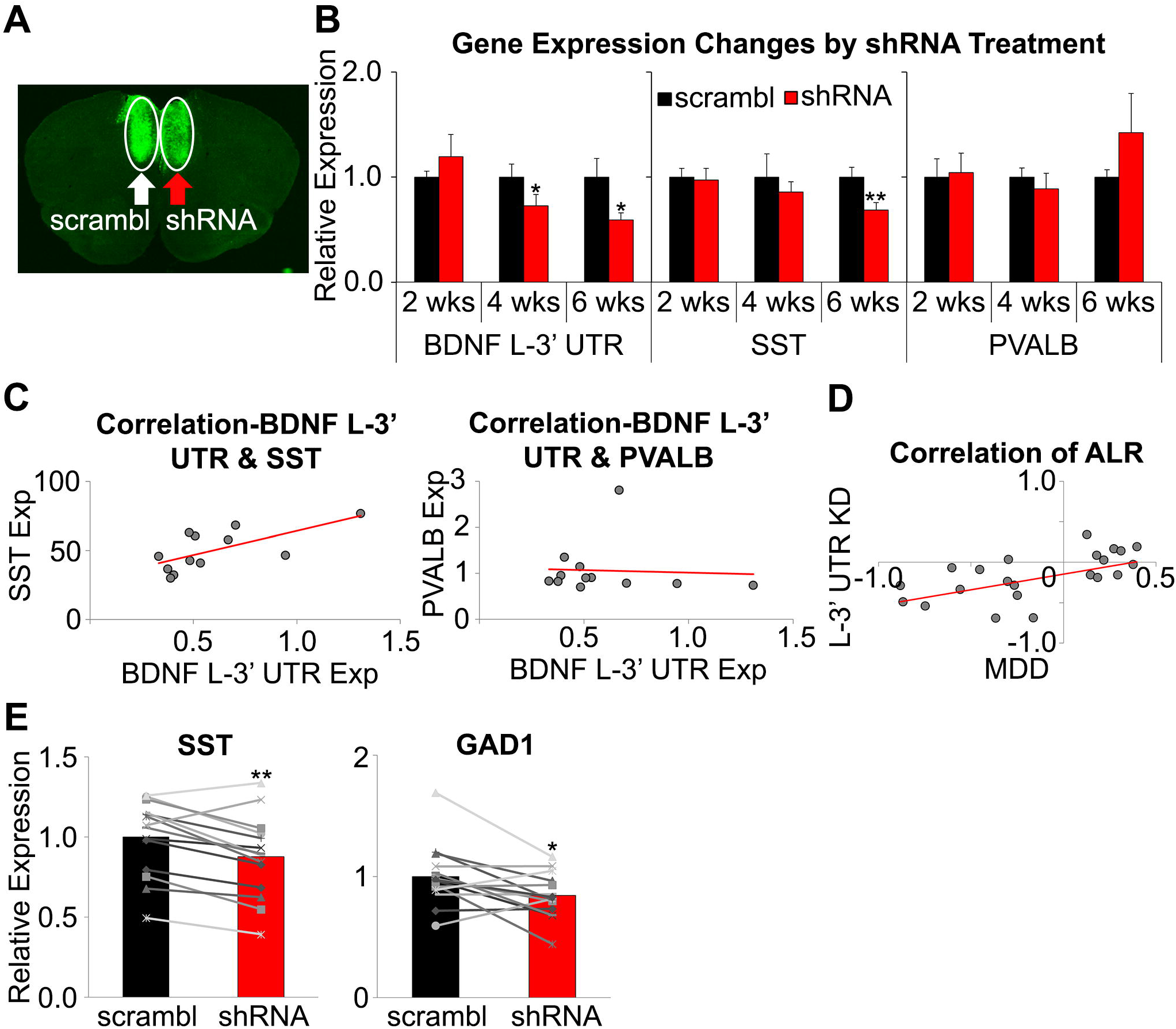
*Bdnf* L-3’-UTR knockdown-related gene expression changes. (A) Representative image of GFP expression following AAV injection in mPFC. (B) Expression changes in core genes at 2, 4, 6 weeks after surgery. Changes in *Bdnf* L-3’-UTR expression were observed from week 4, and *Sst* expression change was observed only in week 6. (C) Correlation between expression of *Bdnf* L-3’-UTR and *Sst* (left) or *Pvalb* (right) level at post-surgery week 6. Significant correlation to *Bdnf* L-3’-UTR level was observed in *Sst*, but not in *Pvalb* (n=6/group at each time point). (D) Similarity between MDD-and *Bdnf* L-3’-UTR knockdown-induced gene expression changes (ALR: average log ratio (MDD/control,BDNF L-3’-UTR KD/control)). (E) *Sst* and *Gad1* in *Sst*+ cells were significantly downregulated by shRNA treatment (n=14/group; *p<0.05, **p<0.01, ***p<0.001). Data are represented as mean ± SEM.

To validate the translational value of our findings, we compared shRNA-induced changes of *Bdnf*-, synaptic function-related genes in mice to expression changes observed in MDD subjects compared to controls. We found a significant correlation (r=0.45, p=0.009; Figure 5D, Table S4), demonstrating that dendritic *Bdnf* knockdown in mice replicated gene expression changes observed in human MDD.

Finally, we collected *Sst*-or *Pvalb*-expressing cells from AAV-infused mPFC using laser microdissection to investigate whether *Bdnf* L-3’-UTR knockdown induced gene expression changes in a cell type-specific manner. Expression of *Sst* and *Gad1* mRNAs was significantly reduced in *Sst*-positive cells by shRNA treatment (p=0.001 and p=0.014, respectively; Figure 5E), but *Pvalb* and *Gad1* remained unchanged in *Pvalb*-positive interneurons (p=0.478 and p=0.217, respectively; Figure S6F-G).

In summary, *Bdnf* L-3’-UTR knockdown recapitulated MDD-and chronic stress-associated molecular, structural, behavioral phenotypes. Our observations that *Sst* reduction followed dendritic *Bdnf* knockdown, and that shRNA-treated mice showed transcriptome profiles similar to MDD suggest that MDD-related dendritic *Bdnf* downregulation is causally implicated in MDD-related GABAergic gene changes and associated symptoms (Figure 6).

**Figure 6.**
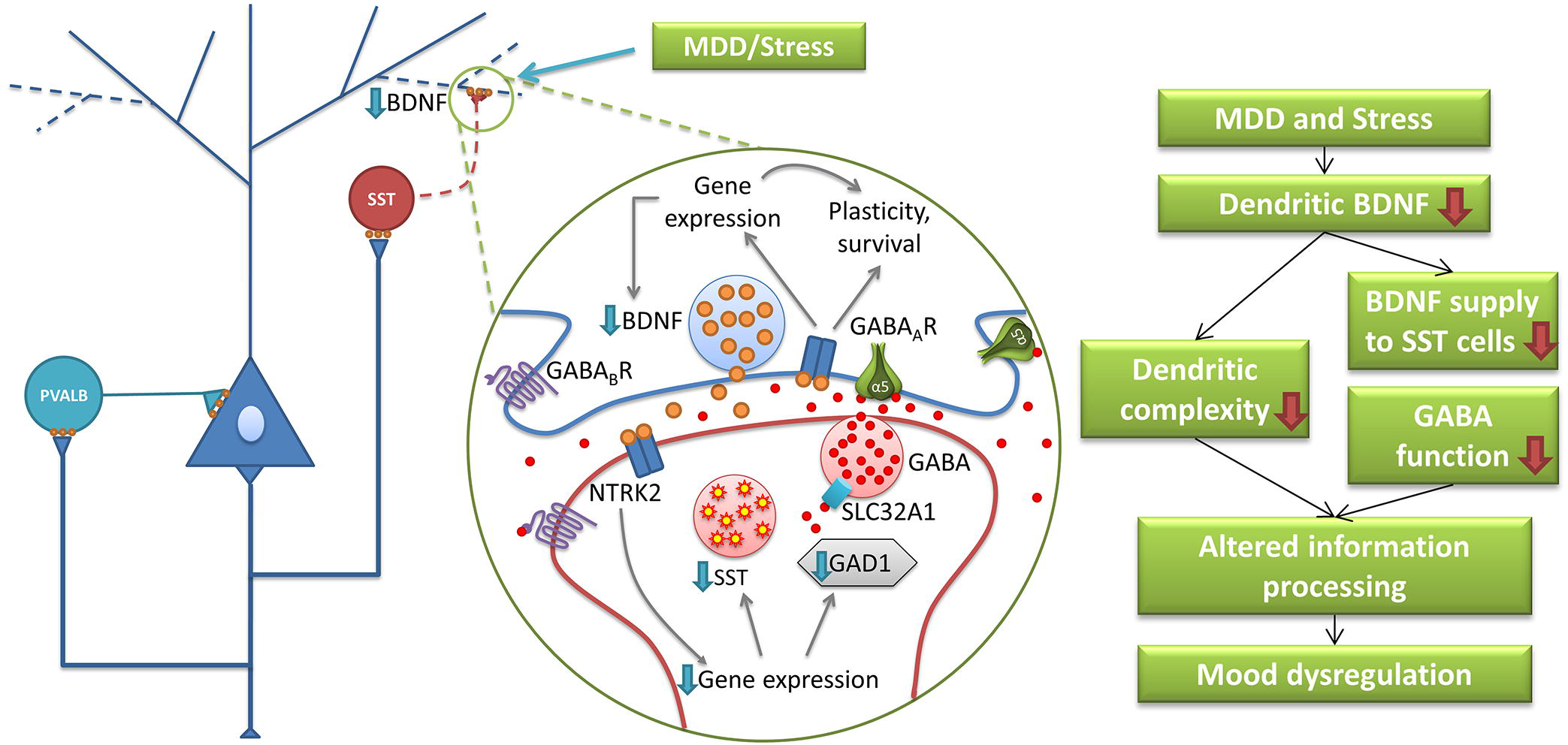
Model of BDNF role in MDD-and stress-related molecular and behavioral phenotypes. Results from the current study suggest a sequence of events associated with MDD-related brain changes. First, prolonged stress induces selective deficit of dendritic BDNF indicated by low BDNF L-3’-UTR expression in the PFC. Second, reduced BDNF supply to dendritic compartments and GABA interneurons targeting distal dendrites of pyramidal cells leads to impaired function of dendritic-targeting interneurons (e.g. downregulation of SST and GABA synthesizing enzyme), disrupted dendritic integrity, altered filtering and processing of excitatory input onto pyramidal cell dendrites, and finally, mood dysregulation.

## DISCUSSION

MDD is a devastating disease characterized by low mood, anhedonia, and cognitive deficits. Studies suggest that impaired neural plasticity and GABAergic function are associated with MDD, however, the mechanism underlying these two phenomena has not been clearly defined. In the current study, we report a selective downregulation of dendritic *BDNF* mRNA in the dlPFC of human MDD subjects and a parallel downregulation of dendritic-targeting interneuron markers without affecting a perisomatic-targeting interneuron marker. Strong and selective correlations between *BDNF* L-3’-UTR and inhibitory synaptic genes were observed, suggesting that an MDD-related decrease in dendritic *BDNF* transcripts contributed to impairment of dendritic-targeting interneurons. A selective deficit of *Bdnf* L-3’-UTR was also observed in the mPFC of mice exposed to UCMS. Although *Sst* expression did not show significant group differences, it was positively correlated with *Bdnf* L-3’-UTR level and negatively to behavioral emotionality. The fact that *Sst* downregulation did not reach statistical significance may relate to differences in timescale; human subjects had suffered from MDD for months to years, whereas mice were exposed to mild stress for only seven weeks. Knockdown of *Bdnf* L-3’-UTR replicated MDD-like structural, behavioral and molecular phenotypes. We observed a trend for elevated anxiety-/depression-like behaviors in shRNA treated mice at baseline, and a significant elevation after UCMS in shRNA-treated animals compared to controls. Following shRNA-induced *Bdnf* knockdown, *Sst* expression was reduced and showed positive correlation to *Bdnf* L-3’-UTR level. These progressive molecular changes were consistent with non-significant group differences in baseline emotionality at post-surgery week 2-3 and with the earlier UCMS-induced fur degradation in shRNA-treated mice. Moreover, cell-specific analyses suggest that *Sst*+ cells are more vulnerable than *Pvalb*+ interneurons to low dendritic *Bdnf* as revealed by reduced *Gad1* expression. Notably, although the mouse studies were inspired by findings in human postmortem brains, species differences may limit their interpretability. Whilst not all of the MDD-associated gene expression changes are recapitulated, we observed a positive correlation between *Bdnf* L-3’-UTR shRNA-induced gene expression changes in mice and those observed in human MDD brain samples. Hence the collective results suggest a novel mechanism underlying the pathophysiology of MDD, linking local neurotrophic support, pyramidal cell structure and dendritic inhibition (Figure 6).

### Contribution of low dendritic *BDNF* to impaired structural integrity and function in the dlPFC of MDD subjects

Previous studies showed that about 50% of *Bdnf* transcripts had L-3’-UTR (31,45). Knockdown of *Bdnf* L-3’-UTR in mice was sufficient to induce stress-and MDD-associated morphological changes in cortical neurons. In agreement with a human postmortem study reporting shortened dendritic processes without changes in the number of processes in basilar dendrites of pyramidal neurons in dlPFC of subjects with psychiatric illnesses including MDD (46) and with animal studies showing stress-induced structural changes mainly occurring on the distal portion of apical dendrites (4,19,20,47,48), shRNA targeting *Bdnf* L-3’-UTR induced retraction of distal dendrites of primary cortical neurons.

Animal studies show that dendritic *Bdnf* transcripts are closely linked to stress-related mental illness (33-35). Human genetic studies also support the idea that the dendritic action of *BDNF* is essential to mood regulation. The most common single nucleotide polymorphism in human *BDNF*, resulting in Valine to Methionine substitution at codon 66 (Val66Met), has been implicated in defective fear extinction (49), and increased susceptibility to psychiatric diseases (50). Mice harboring the Met allele exhibit impaired dendritic transport of *Bdnf* mRNA (51,52), reduced activity-dependent *Bdnf* secretion (53), impaired synaptic plasticity in mPFC (54), and elevated anxiety-like behavior which is not normalized by fluoxetine (55). Our finding that emotionality is significantly correlated to *Bdnf* L-3’-UTR expression provides evidence that dendritic *Bdnf* is important for the function of mood regulatory systems.

Then how can the loss of dendritic *BDNF* contribute to functional abnormalities? First, defects in activity-dependent *BDNF* supply might play a role. A recent study found that over 2,000 transcripts are localized in dendrites and/or axons, maybe for faster on-demand protein supply by local synaptic translation (32). Not only does L-3’-UTR+ *Bdnf* mRNA preferentially exist in dendrites (31), it stays dormant in resting state, yet binds to a polyribosome responding to neuronal activity (45). Given that MDD may be conceived as a disease of maladaptive processes, impaired activity-dependent neuroplasticity may underlie the inability to cope with stress. A second possible mechanism could be through reduced GABAergic transmission. Knockdown of dendritic *Bdnf* mRNA and/or blockade of dendritic transport downregulate GABAergic receptors and GABA synthesizing enzymes (56,57). In addition, *Sst* acts as inhibitory neuropeptide and *Sst* knockout is sufficient to induce depressive-/anxiety-related behavior in mice (16). These data imply that the depressive-like behavioral outcome may derive at least in part from low inhibitory control.

### Close link between pyramidal cell dendritic *BDNF* transcripts and dendritic-targeting interneurons and its implication for the pathology of MDD

*SST* and *PVALB* are expressed in distinct classes of cortical interneurons. Multiple lines of evidence imply that *BDNF*-*NTRK2* signaling plays a crucial role in interneuron development and maintenance of GABAergic function. Our studies have consistently found that low *BDNF* function is closely linked to reduced expression of GABA-related genes observed in the brains of MDD patients (11,14,24). As *NTRK2* is predominantly expressed in *PVALB*+ interneurons compared to other interneuron populations (21,22), higher *BDNF* dependency of *SST* than *PVALB* raised questions about underlying mechanisms. Our current study suggests that selectivity of MDD/stress-induced effects may be achieved through reduced dendritic *BDNF* function, leading to altered neuronal plasticity, behavior and impaired function of dendritic-targeting interneurons.

To date, it is not clearly defined how dysfunction of *SST*+ interneurons contribute to the pathophysiology of mood disorders (58). In a previous study, we reported that acute inactivation of *SST* neurons in mPFC can increase anxiety-and depressive-like behavior (59). The mPFC, the target of our study, has excitatory projections to amygdala to produce fear/stress response (60). Hence low *BDNF* L-3’-UTR-mediated dysfunction of *SST* neurons may result in increased mPFC pyramidal neuron baseline activity leading to amygdala hyperactivity. In addition, distal dendrites mainly receive inputs from other cortical areas and regulation of dendritic excitability is crucial for functional computations (reviewed in Larkum (61)). Considering that dendritic excitation mainly occurs in actively-behaving animals, one can assume that proper control of dendritic activity is crucial for regulation of active behavior. For instance, *Sst*+ interneurons in barrel cortex are inactivated during active whisking which leads to increased excitability for the sensorimotor stimuli (62). During contextual fear learning, *Sst*+ interneurons in the basolateral amygdala are inactivated by multisensory environmental context, whereas an aversive event inactivates both *Pvalb*+ and *Sst*+ interneurons (63,64). Hence, loss of dendritic inhibition may result in increased signal–to-noise ratio and failure to discriminate different stimuli (58).

Combining current knowledge and our experimental results, we propose a biological model linking low *BDNF*, structural changes and GABAergic dysfunction, three phenomena commonly observed in the PFC of MDD subjects and stressed animals (Figure 6). MDD and stress may disturb dendritic *BDNF*, which leads to low neurotrophic supply to dendrites and dendritic targeting interneurons. Reduced *BDNF* signaling may contribute to decreased dendritic complexity of pyramidal neurons and low inhibitory function via reduced *GAD* (i.e. low GABA production) and *SST* expression in *SST*+ cells, finally culminating in altered information processing by cortical microcircuits, leading to behavioral symptoms of MDD.

In conclusion, our results contribute to our understanding of *BDNF* function under control and disease-states, and provide evidence that MDD-related downregulation of dendritic *BDNF* contributes to selective impairment of dendritic-targeting interneurons in MDD subjects, and that maintaining the integrity of the *BDNF-NTRK2* system is critical for mood regulation.

### Limitations

First of all, we did not assess BDNF protein level in the mouse brains after BDNF L-3’-UTR knockdown because of limited amount of tissue. Technical advances, especially protein quantification method, might allow addressing this question. Second, we utilized postmortem human brains which do not allow examining dynamic molecular changes induced by disease. Therefore, our findings could be the results of the disease, not the cause. Although UCMS paradigm is known to reproduce phenotypes resembling human MDD (65), it is a model of chronic stress-induced pathologies, rather than MDD itself. Third, we performed animal experiments with male mice. Given that SST expression changes is greater in female MDD subjects than in males, we expect that female mice would show greater, significant SST changes after UCMS.

## ACKNOWLEDGEMENTS

This work was supported by National Institute of Mental Health (NIMH) MH077159 (ES).

## REFERENCES

1. Kupfer DJ, Frank E, Phillips ML (2012): Major depressive disorder: new clinical, neurobiological, and treatment perspectives. Lancet. 379:1045–1055.

2. McEwen BS, Morrison JH (2013): The Brain on Stress: Vulnerability and Plasticity of the Prefrontal Cortex over the Life Course. Neuron. 79:16–29.

3. Drevets WC (2000): Functional anatomical abnormalities in limbic and prefrontal cortical structures in major depression. Prog Brain Res. 126:413–431.

4. Liston C, Miller MM, Goldwater DS, Radley JJ, Rocher AB, Hof PR, et al. (2006): Stress-induced alterations in prefrontal cortical dendritic morphology predict selective impairments in perceptual attentional set-shifting. J Neurosci. 26:7870–7874.

5. Tan HY, Callicott JH, Weinberger DR (2007): Dysfunctional and compensatory prefrontal cortical systems, genes and the pathogenesis of schizophrenia. Cereb Cortex. 17 Suppl 1:i171–181.

6. Kang HJ, Voleti B, Hajszan T, Rajkowska G, Stockmeier CA, Licznerski P, et al. (2012): Decreased expression of synapse-related genes and loss of synapses in major depressive disorder. Nat Med. 18:1413–1417.

7. Rajkowska G (2000): Postmortem studies in mood disorders indicate altered numbers of neurons and glial cells. Biol Psychiatry. 48:766–777.

8. Levinson AJ, Fitzgerald PB, Favalli G, Blumberger DM, Daigle M, Daskalakis ZJ (2010): Evidence of cortical inhibitory deficits in major depressive disorder. Biol Psychiatry. 67:458–464.

9. Sanacora G, Mason GF, Rothman DL, Behar KL, Hyder F, Petroff OAC, et al. (1999): Reduced cortical gamma-aminobutyric acid levels in depressed patients determined by proton magnetic resonance spectroscopy. Arch Gen Psychiat. 56:1043–1047.

10. Sanacora G, Gueorguieva R, Epperson CN, Wu YT, Appel M, Rothman DL, et al. (2004): Subtype-specific alterations of gamma-aminobutyric acid and glutamate in patients with major depression. Arch Gen Psychiat. 61:705–713.

11. Guilloux JP, Douillard-Guilloux G, Kota R, Wang X, Gardier AM, Martinowich K, et al. (2012): Molecular evidence for BDNF- and GABA-related dysfunctions in the amygdala of female subjects with major depression. Mol Psychiatry. 17:1130–1142.

12. Sibille E, Morris HM, Kota RS, Lewis DA (2011): GABA-related transcripts in the dorsolateral prefrontal cortex in mood disorders. Int J Neuropsychopharmacol. 14:721–734.

13. Tripp A, Kota RS, Lewis DA, Sibille E (2011): Reduced somatostatin in subgenual anterior cingulate cortex in major depression. Neurobiol Dis. 42:116–124.

14. Tripp A, Oh H, Guilloux JP, Martinowich K, Lewis DA, Sibille E (2012): Brain-derived neurotrophic factor signaling and subgenual anterior cingulate cortex dysfunction in major depressive disorder. Am J Psychiatry. 169:1194–1202.

15. Lin LC, Sibille E (2013): Reduced brain somatostatin in mood disorders: a common pathophysiological substrate and drug target? Front Pharmacol. 4:110.

16. Lin LC, Sibille E (2015): Somatostatin, neuronal vulnerability and behavioral emotionality. Mol Psychiatry. 20:377–387.

17. Duman RS, Monteggia LM (2006): A neurotrophic model for stress-related mood disorders. Biol Psychiatry. 59:1116–1127.

18. Dunham JS, Deakin JF, Miyajima F, Payton A, Toro CT (2009): Expression of hippocampal brain-derived neurotrophic factor and its receptors in Stanley consortium brains. Journal of psychiatric research. 43:1175–1184.

19. Radley JJ, Rocher AB, Rodriguez A, Ehlenberger DB, Dammann M, McEwen BS, et al. (2008): Repeated stress alters dendritic spine morphology in the rat medial prefrontal cortex. J Comp Neurol. 507:1141–1150.

20. Radley JJ, Sisti HM, Hao J, Rocher AB, McCall T, Hof PR, et al. (2004): Chronic behavioral stress induces apical dendritic reorganization in pyramidal neurons of the medial prefrontal cortex. Neuroscience. 125:1–6.

21. Cellerino A, Maffei L, Domenici L (1996): The distribution of brain-derived neurotrophic factor and its receptor trkB in parvalbumin-containing neurons of the rat visual cortex. The European journal of neuroscience. 8:1190–1197.

22. Gorba T, Wahle P (1999): Expression of TrkB and TrkC but not BDNF mRNA in neurochemically identified interneurons in rat visual cortex in vivo and in organotypic cultures. The European journal of neuroscience. 11:1179–1190.

23. Kohara K, Yasuda H, Huang Y, Adachi N, Sohya K, Tsumoto T (2007): A local reduction in cortical GABAergic synapses after a loss of endogenous brain-derived neurotrophic factor, as revealed by single-cell gene knock-out method. J Neurosci. 27:7234–7244.

24. Glorioso C, Sabatini M, Unger T, Hashimoto T, Monteggia LM, Lewis DA, et al. (2006): Specificity and timing of neocortical transcriptome changes in response to BDNF gene ablation during embryogenesis or adulthood. Mol Psychiatry. 11:633–648.

25. Wang L, Chang X, She L, Xu D, Huang W, Poo MM (2015): Autocrine action of BDNF on dendrite development of adult-born hippocampal neurons. J Neurosci. 35:8384–8393.

26. Tanaka J, Horiike Y, Matsuzaki M, Miyazaki T, Ellis-Davies GC, Kasai H (2008): Protein synthesis and neurotrophin-dependent structural plasticity of single dendritic spines. Science. 319:1683–1687.

27. Ohba S, Ikeda T, Ikegaya Y, Nishiyama N, Matsuki N, Yamada MK (2005): BDNF locally potentiates GABAergic presynaptic machineries: target-selective circuit inhibition. Cereb Cortex. 15:291–298.

28. Ali AB, Thomson AM (2008): Synaptic alpha 5 subunit-containing GABAA receptors mediate IPSPs elicited by dendrite-preferring cells in rat neocortex. Cereb Cortex. 18:1260–1271.

29. Oh H, Lewis DA, Sibille E (2016): The Role of BDNF in Age-Dependent Changes of Excitatory and Inhibitory Synaptic Markers in the Human Prefrontal Cortex. Neuropsychopharmacology. 41:3080–3091.

30. Baj G, Leone E, Chao MV, Tongiorgi E (2011): Spatial segregation of BDNF transcripts enables BDNF to differentially shape distinct dendritic compartments. Proc Natl Acad Sci U S A. 108:16813–16818.

31. An JJ, Gharami K, Liao GY, Woo NH, Lau AG, Vanevski F, et al. (2008): Distinct role of long 3’ UTR BDNF mRNA in spine morphology and synaptic plasticity in hippocampal neurons. Cell. 134:175–187.

32. Cajigas IJ, Tushev G, Will TJ, tom Dieck S, Fuerst N, Schuman EM (2012): The local transcriptome in the synaptic neuropil revealed by deep sequencing and high-resolution imaging. Neuron. 74:453–466.

33. Luoni A, Berry A, Calabrese F, Capoccia S, Bellisario V, Gass P, et al. (2014): Delayed BDNF alterations in the prefrontal cortex of rats exposed to prenatal stress: preventive effect of lurasidone treatment during adolescence. European neuropsychopharmacology : the journal of the European College of Neuropsychopharmacology. 24:986–995.

34. Luoni A, Macchi F, Papp M, Molteni R, Riva MA (2015): Lurasidone Exerts Antidepressant Properties in the Chronic Mild Stress Model through the Regulation of Synaptic and Neuroplastic Mechanisms in the Rat Prefrontal Cortex. Int J Neuropsychoph. 18.

35. Berry A, Panetta P, Luoni A, Bellisario V, Capoccia S, Riva MA, et al. (2015): Decreased Bdnf expression and reduced social behavior in periadolescent rats following prenatal stress. Developmental psychobiology. 57:365–373.

36. Baj G, D’Alessandro V, Musazzi L, Mallei A, Sartori CR, Sciancalepore M, et al. (2012): Physical Exercise and Antidepressants Enhance BDNF Targeting in Hippocampal CA3 Dendrites: Further Evidence of a Spatial Code for BDNF Splice Variants. Neuropsychopharmacology. 37:1600–1611.

37. Orefice LL, Waterhouse EG, Partridge JG, Lalchandani RR, Vicini S, Xu B (2013): Distinct roles for somatically and dendritically synthesized brain-derived neurotrophic factor in morphogenesis of dendritic spines. J Neurosci. 33:11618–11632.

38. Glantz LA, Austin MC, Lewis DA (2000): Normal cellular levels of synaptophysin mRNA expression in the prefrontal cortex of subjects with schizophrenia. Biol Psychiatry. 48:389–397.

39. Glantz LA, Lewis DA (1997): Reduction of synaptophysin immunoreactivity in the prefrontal cortex of subjects with schizophrenia. Regional and diagnostic specificity. Arch GenPsychiatry. 54:660–669.

40. Sweet RA, Bergen SE, Sun Z, Sampson AR, Pierri JN, Lewis DA (2004): Pyramidal cell size reduction in schizophrenia: evidence for involvement of auditory feedforward circuits. BiolPsychiatry. 55:1128–1137.

41. Sibille E, Arango V, Galfalvy HC, Pavlidis P, Erraji-BenChekroun L, Ellis SP, et al. (2004): Gene expression profiling of depression and suicide in human prefrontal cortex. Neuropsychopharmacology. 29:351–361.

42. Guilloux JP, Seney M, Edgar N, Sibille E (2011): Integrated behavioral z-scoring increases the sensitivity and reliability of behavioral phenotyping in mice: Relevance to emotionality and sex. J Neurosci Methods. 197:21–31.

43. Waterhouse EG, Xu BJ (2009): New insights into the role of brain-derived neurotrophic factor in synaptic plasticity. Mol Cell Neurosci. 42:81–89.

44. Vicario A, Colliva A, Ratti A, Davidovic L, Baj G, Gricman L, et al. (2015): Dendritic targeting of short and long 3’ UTR BDNF mRNA is regulated by BDNF or NT-3 and distinct sets of RNA-binding proteins. Front Mol Neurosci. 8:62.

45. Lau AG, Irier HA, Gu JP, Tian DH, Ku L, Liu GL, et al. (2010): Distinct 3 ’ UTRs differentially regulate activity-dependent translation of brain-derived neurotrophic factor (BDNF). P Natl Acad Sci USA. 107:15945–15950.

46. Glantz LA, Lewis DA (2000): Decreased dendritic spine density on prefrontal cortical pyramidal neurons in schizophrenia. Arch Gen Psychiatry. 57:65–73.

47. Bloss EB, Janssen WG, McEwen BS, Morrison JH (2010): Interactive effects of stress and aging on structural plasticity in the prefrontal cortex. J Neurosci. 30:6726–6731.

48. Cook SC, Wellman CL (2004): Chronic stress alters dendritic morphology in rat medial prefrontal cortex. J Neurobiol. 60:236–248.

49. Soliman F, Glatt CE, Bath KG, Levita L, Jones RM, Pattwell SS, et al. (2010): A Genetic Variant BDNF Polymorphism Alters Extinction Learning in Both Mouse and Human. Science. 327:863–866.

50. Gatt JM, Nemeroff CB, Dobson-Stone C, Paul RH, Bryant RA, Schofield PR, et al. (2009): Interactions between BDNF Val66Met polymorphism and early life stress predict brain and arousal pathways to syndromal depression and anxiety. Mol Psychiatr. 14:681–695.

51. Chiaruttini C, Vicario A, Li Z, Baj G, Braiuca P, Wu Y, et al. (2009): Dendritic trafficking of BDNF mRNA is mediated by translin and blocked by the G196A (Val66Met) mutation. Proc Natl Acad Sci U S A. 106:16481–16486.

52. Mallei A, Baj G, Ieraci A, Corna S, Musazzi L, Lee FS, et al. (2015): Expression and Dendritic Trafficking of BDNF-6 Splice Variant are Impaired in Knock-In Mice Carrying Human BDNF Val66Met Polymorphism. Int J Neuropsychopharmacol.

53. Egan MF, Kojima M, Callicott JH, Goldberg TE, Kolachana BS, Bertolino A, et al. (2003): The BDNF val66met polymorphism affects activity-dependent secretion of BDNF and human memory and hippocampal function. Cell. 112:257–269.

54. Pattwell SS, Bath KG, Perez-Castro R, Lee FS, Chao MV, Ninan I (2012): The BDNF Val66Met polymorphism impairs synaptic transmission and plasticity in the infralimbic medial prefrontal cortex. J Neurosci. 32:2410–2421.

55. Chen ZY, Jing D, Bath KG, Ieraci A, Khan T, Siao CJ, et al. (2006): Genetic variant BDNF (Val66Met) polymorphism alters anxiety-related behavior. Science. 314:140–143.

56. Stein JM, Bergman W, Fang Y, Davison L, Brensinger C, Robinson MB, et al. (2006): Behavioral and neurochemical alterations in mice lacking the RNA-binding protein translin. J Neurosci. 26:2184–2196.

57. Waterhouse EG, An JJ, Orefice LL, Baydyuk M, Liao GY, Zheng K, et al. (2012): BDNF Promotes Differentiation and Maturation of Adult-born Neurons through GABAergic Transmission. Journal of Neuroscience. 32:14318–14330.

58. Fee C, Banasr M, Sibille E (2017): Somatostatin-Positive Gamma-Aminobutyric Acid Interneuron Deficits in Depression: Cortical Microcircuit and Therapeutic Perspectives. Biol Psychiatry.

59. Soumier A, Sibille E (2014): Opposing effects of acute versus chronic blockade of frontal cortex somatostatin-positive inhibitory neurons on behavioral emotionality in mice. Neuropsychopharmacology. 39:2252.

60. Vidal-Gonzalez I, Vidal-Gonzalez B, Rauch SL, Quirk GJ (2006): Microstimulation reveals opposing influences of prelimbic and infralimbic cortex on the expression of conditioned fear. Learning & memory. 13:728–733.

61. Larkum M (2013): A cellular mechanism for cortical associations: an organizing principle for the cerebral cortex. Trends Neurosci. 36:141–151.

62. Gentet LJ, Kremer Y, Taniguchi H, Huang ZJ, Staiger JF, Petersen CC (2012): Unique functional properties of somatostatin-expressing GABAergic neurons in mouse barrel cortex. Nat Neurosci. 15:607–612.

63. Tovote P, Fadok JP, Luthi A (2015): Neuronal circuits for fear and anxiety. Nat Rev Neurosci. 16:317–331.

64. Wolff SB, Grundemann J, Tovote P, Krabbe S, Jacobson GA, Muller C, et al. (2014): Amygdala interneuron subtypes control fear learning through disinhibition. Nature. 509:453–458.

65. Nollet M, Guisquet AML, Belzung C Models of depression: unpredictable chronic mild stress in mice. Current protocols in pharmacology.5.65. 61–65.65. 17.

66. Franklin KBJ, Paxinos G (2008): The Mouse Brain in Stereotaxic Coordinates. Academic Press.

